# Reduced graphene oxide membrane as supporting film for high-resolution cryo-EM

**DOI:** 10.1101/2021.04.15.439953

**Authors:** Nan Liu, Liming Zheng, Jie Xu, Jia Wang, Cuixia Hu, Jun Lan, Xing Zhang, Jincan Zhang, Kui Xu, Hang Cheng, Zi Yang, Xin Gao, Xinquan Wang, Hailin Peng, Yanan Chen, Hong-Wei Wang

## Abstract

Although single-particle cryogenic electron microscopy (cryo-EM) has been applied extensively for elucidating many crucial biological mechanisms at the molecular level, this technique still faces critical challenges, the major one of which is to prepare the high-quality cryo-EM specimen. Aiming to achieve a more reproducible and efficient cryo-EM specimen preparation, novel supporting films including graphene-based two-dimensional materials have been explored in recent years. Here we report a robust and simple method to fabricate EM grids coated with single- or few-layer reduced graphene oxide (RGO) membrane in large batch for high-resolution cryo-EM structural determination. The RGO membrane has decreased interlayer space and enhanced electrical conductivity in comparison to regular graphene oxide (GO) membrane. Moreover, we found that the RGO supporting film exhibited nice particle-absorption ability, thus avoiding the air-water interface problem. More importantly, we found that the RGO supporting film is particularly useful in cryo-EM reconstruction of sub-100 kDa biomolecules at near-atomic resolution, as exemplified by the study of RBD-ACE2 complex and other small protein molecules. We envision that the RGO membranes can be used as a robust graphene-based supporting film in cryo-EM specimen preparation.

## Introduction

Single-particle cryo-EM, without the requirement of crystallization, has become a major method to solve structures of biological macromolecules at near-atomic resolution benefiting from its recent advances in both software and hardware (Cheng, 2015; Cheng, 2018; Li et al., 2013; Scheres, 2012; Wu et al., 2016). The more recent breakthroughs even pushed the resolution to genuine atomic level (Nakane et al., 2020; Yip et al., 2020). Furthermore, the method enables the analysis of heterogeneity to solve multiple conformers from one single cryo-EM dataset (Scheres, 2012). While cryo-EM has become one of the most powerful methods in structural biology, a major bottleneck limiting the method’s general application is the poor reproducibility and controllability to prepare cryo-EM specimen (Armstrong et al., 2020; Glaeser, 2016, 2018). The conventional and most popularly used cryo-EM specimen preparation procedure is to apply a droplet of solution containing target macromolecules onto an EM grid coated with holey carbon film, to blot the grid with filter paper and then to plunge-freeze the grid into liquid nitrogen temperature to form a very thin vitreous ice embedding the macromolecules (Dubochet et al., 1982; Grassucci et al., 2007). During the blotting process to form thin liquid film, biomolecular particles with protein components tend to be absorbed onto the air-water interface, causing denaturation and preferred orientation (Glaeser, 2018; Noble et al., 2018). The adsorption of molecules to the air-water interface was hypothesized to be the main reason of low yield in cryo-EM specimen preparation (D’Imprima et al., 2019; Noble et al., 2018). To overcome this problem, thin films of various materials have been developed to support the biological macromolecules over holey grids. The most frequently used thin film is continuous amorphous carbon, which contributes to strong background noise, aggravate charging effect and beam-induced motion due to its poor electrical conductivity when imaged under electron microscopy (Russo and Passmore, 2014a, c; Zheng et al., 2020).

Graphene (Geim and Novoselov, 2007), a two-dimensional nano-material with the superior properties of ultrahigh electrical/thermal conductivity (Balandin et al., 2008; Chen et al., 2008), mechanical strength (Lee et al., 2008) and low background noise (Russo and Passmore, 2014a), is considered as an ideal supporting film for cryo-EM specimen preparation. Graphene membrane as well as its derivatives, such as functionalized graphene (D’Imprima et al., 2019; Han et al., 2020; Liu et al., 2019; Naydenova et al., 2019; Russo and Passmore, 2014b), have been reported to help successfully determining high-resolution reconstruction of multiple macromolecular structures by cryo-EM. The synthesis of single-crystalline graphene with large area, however, due to its high technical and resource demand (Lin et al., 2016), is difficult to be established in regular laboratories focusing on biology. Moreover, the preparation of CVD-prepared or commercially available graphene onto EM grids normally involves many chemical reagents and is hard to avoid contaminations (Han et al., 2020; Regan et al., 2010; Zhang et al., 2017). On the other hand, graphene oxide (GO) containing plenty of functional groups like carboxyl or epoxy (Pantelic et al., 2010) can be easily generated by oxidizing graphite, therefore has been explored as supporting film for cryo-EM specimen preparation (Benjamin et al., 2016; Palovcak et al., 2018; Pantelic et al., 2010; Wang et al., 2020; Wilson et al., 2009). However, GO membrane is electrical insulative (Jung et al., 2008; Wang et al., 2018), which may cause charging accumulation in the irradiated region (Egerton et al., 2004). In addition, the space of interlayer in multiple-layer GO films is larger than that of multi-layer graphene because of the abundant functional groups and encapsulated solvent molecules in the GO films, introducing extra background noise (Moon et al., 2010; Qiu et al., 2015).

Herein, we develop a facile and robust strategy to use reduced graphene oxide (RGO) membrane as supporting substrate for cryo-EM specimen preparation. Compared with GO, RGO contained fewer functional groups with decreased interlayer space and better electric conductivity. Notably, the RGO membrane enabled nice absorption of target biomolecules and high-resolution cryo-EM reconstruction. Several sub-100 kDa biomolecules exhibited nice contrast on RGO membrane, and we successfully solved the structure of SARS-CoV2 RBD-ACE2 complex at 2.8 Å resolution. In our practice, the RGO membrane seemed particularly useful for cryo-EM analysis of relatively small protein molecules in low concentration.

## Results

### Fabrication and characterization of RGO grids

Graphene oxide (GO) could be produced by improved Hummers’ method (Marcano et al., 2010) or commercially purchased, and dispersed in methanol/water solution. The GO layers were firstly coated onto holey-carbon EM grids following the previously reported method (Wang et al., 2020). The grids were then baked in an atmosphere of hydrogen/argon to reduce the GO membranes into RGO (Method, Schema 1 and Figure S1). RGO grids can be simply fabricated in large batch and we can normally prepare several hundred RGO grids at a time.

**Schema 1.**
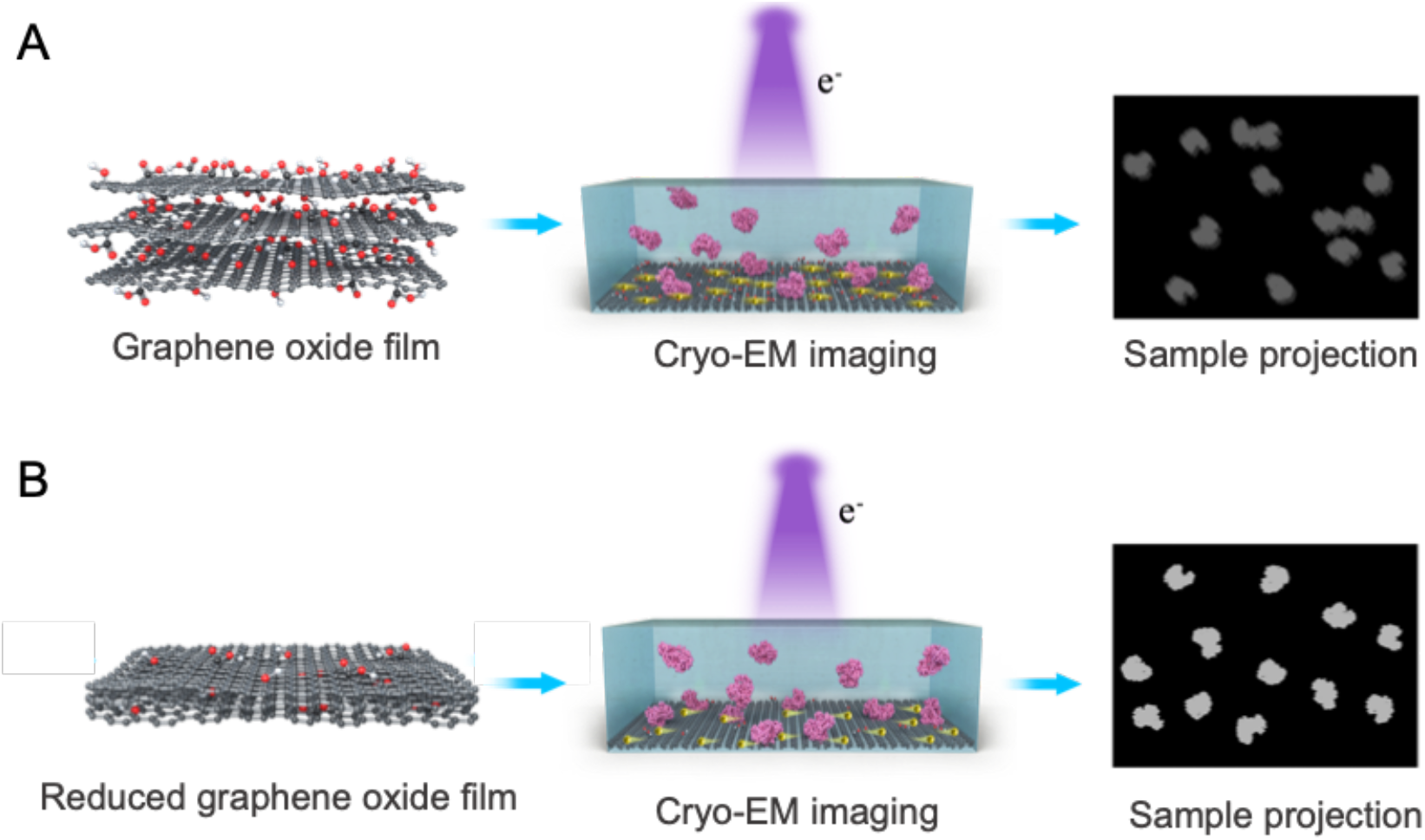
Schematic diagram of graphene oxide (**A**) and reduced graphene oxide (**B**) for single-particle cryo-EM analysis.

We characterized the reduction efficiency of functional groups on GO by X-ray photoelectron spectroscopy (XPS) and found that the O1s peak intensity decreased after reduction treatment (Figure 1A and Figure S2). The atomic ratio (O1s/C1s) of RGO was reduced to 0.16 from 0.38 of GO, consistent with the reported results by chemical graphitization (Moon et al., 2010). We further analyzed the high-resolution spectra of C1s. The signals of C-O (binding energy 286.7 eV) and O-C=O (288.4 eV) bonds which were evident in GO entirely disappeared in RGO, where the sp^2^ carbon of C=C (284.5 eV) and C-C (285.3 eV) dominated the bonding species (Figure 1B and C). Accordingly, the peak area ratio of oxygen functional groups in RGO was down to 0.06, which was significantly decreased in comparison with that (~0.47) of GO. The spectrum analysis demonstrated that a large number of oxygen-containing functional groups on graphene oxide layers had been successfully eliminated by the reduction treatment.

**Figure 1.**
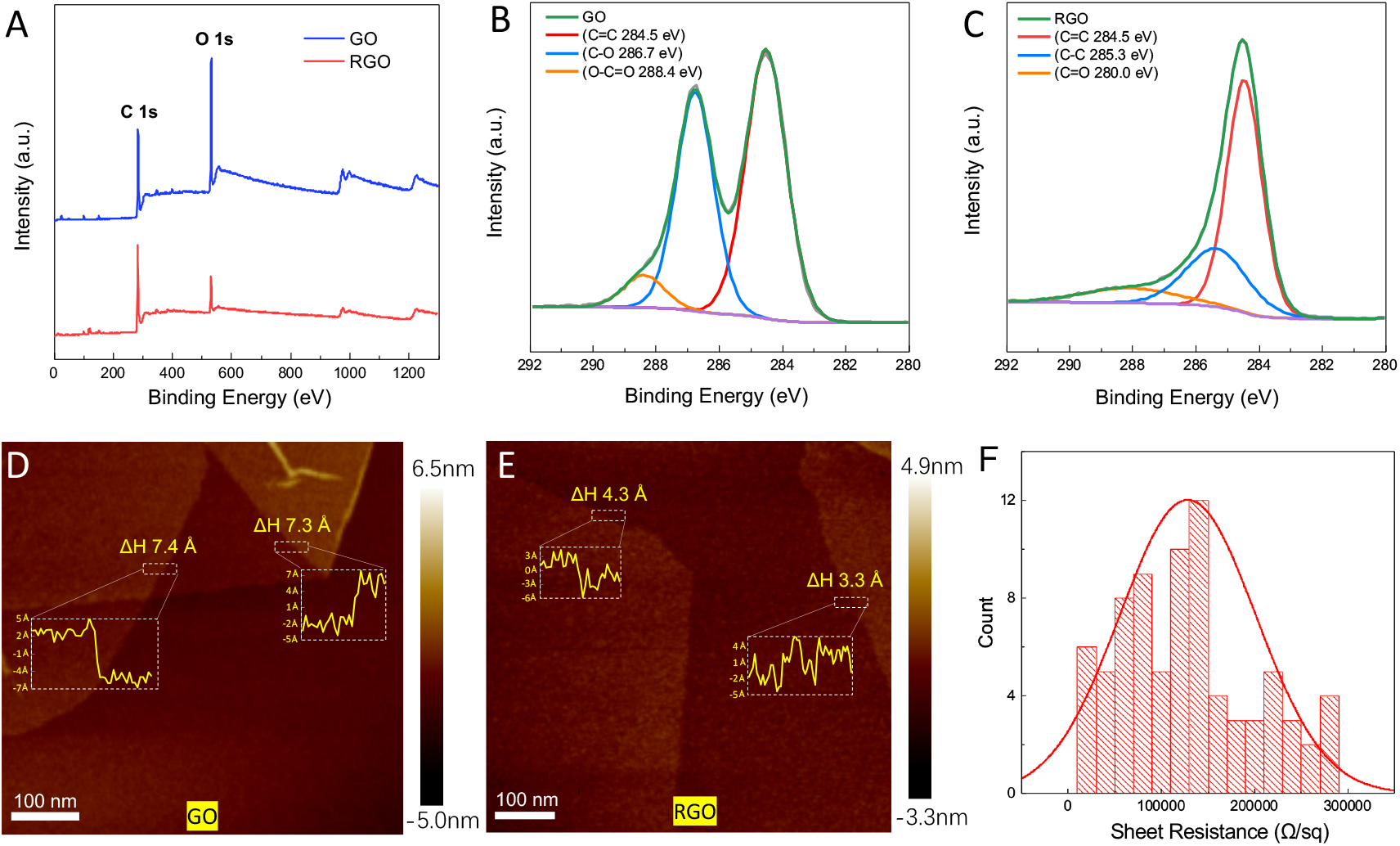
Characterization of reduced graphene oxide membrane. **A**. XPS spectra of graphene oxide (GO) and reduced graphene oxide (RGO). Energy binding peaks assigned to C1s and O1s of 284.4 eV and 532.8 eV were labeled. **B**. High-resolution XPS C1s spectra of graphene oxide. **C**. High-resolution XPS C1s spectra of reduced graphene oxide. **D**. Interlayer space of graphene oxide membrane measured by AFM. The measured regions and corresponding height plots were windowed by rectangles. **E**. Interlayer space of reduced graphene oxide membrane measured by AFM. **F.** Sheet-resistance histogram of reduced graphene oxide membrane.

In order to measure the interlayer distance of GO and RGO membranes, we coated both samples on fresh mica plate and examined them by atomic force microscopy (AFM). The interlayer space can be measured by the height profile of two neighboring layers. The space of RGO was 0.3~0.4 nm (Figure 1E), consistent with previous reports (Lian et al., 2018; Moon et al., 2010), about half of that of GO with over 0.7 nm (Figure 1D) (Buchsteiner et al., 2006; Lian et al., 2018; You et al., 2013). The interlayer space reduction can be explained by two reasons. First, the branching functional groups covalently bound onto GO layers stretch the interlayer space, while the layers of RGO stack more tightly due to the lack of branching functional groups. Second, the GO membranes with active functional groups may attract small molecules like water, further expanding the interlayer space (Lian et al., 2018). The interlayer space of RGO is similar to that of the CVD-prepared multilayer graphene membranes of ~0.35nm, further indicating a similar structure of RGO to graphene. We further tested the electrical conductivity of GO and RGO to see which one is more conductive as the supporting material in EM. We verified that the GO sheet was electrically insulative (Qiu et al., 2015; Wang et al., 2018) and its sheet-resistance exceeded the measuring range in our characterization. In contrast, the sheet-resistance of RGO was dramatically reduced (~10^5^ Ω per square), and the lowest resistance can be 20,000 Ω per square (Figure 1F).

### Characterization of the hydrophilicity and charging effect of RGO membrane

The coverage of RGO membrane on EM grids prepared as described above was more than 90% and free of contamination (Figure 2A, and Figure S3-S4). The reduction procedures barely broke the graphene membrane and kept the layer number of GO as in the initial coating (Figure S3). It was ordinary to find single-layer graphene with good crystallinity covering an entire hole, as indicated by the selected area electron-diffraction pattern (Figure 2A, and Figure S4). We further evaluated the layer number of graphene across holes and found that more than half of the counted holes were covered by single-layer graphene and ~40% were covered by two-layer graphene (Figure 2B), which verified that the majority of the holes were covered by single- or few-layer graphene film, thus generating much less background noise than conventional amorphous carbon supporting film. We tested the hydrophilicity of GO and RGO coated grids, by measuring the water contact angle (WCA) after low-energy plasma treatments. As expected, RGO grids with fewer functional groups exhibited larger WCA compared to GO grids, indicating stronger hydrophobicity. However, their WCAs were rapidly decreased when treated with plasma cleaning (Figure 2C), suitable to be used for cryo-EM specimen preparation.

**Figure 2.**
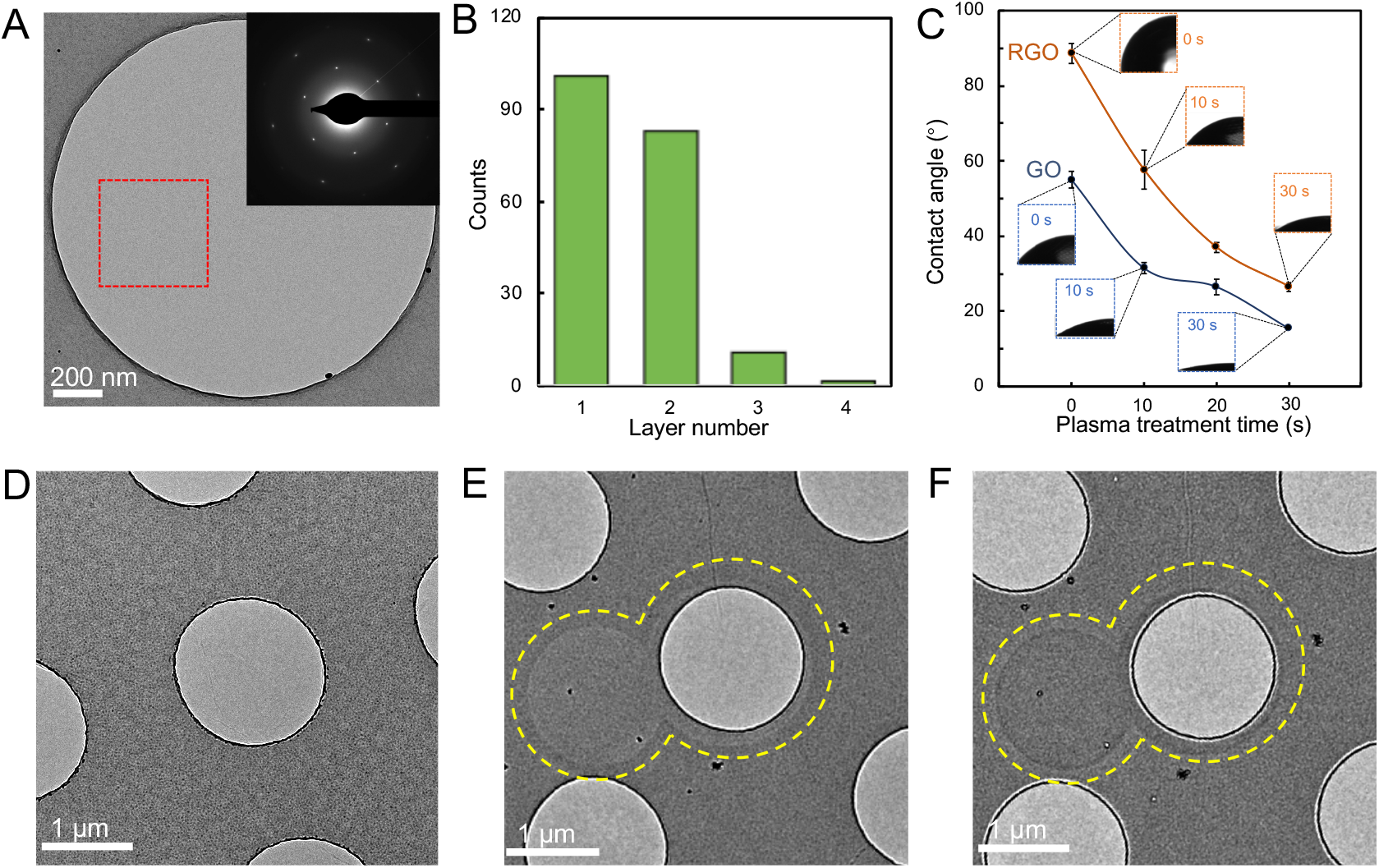
TEM characterization and hydrophilicity of RGO and GO membranes. **A**. A representative TEM image of RGO monolayer covering a hole. The inset was the electron diffraction pattern of the selected region labeled by red square. **B**. Statistics of graphene layer number across holes on the EM grid. **C**. Water contact angles of RGO (orange curve) and GO (dark blue) coated grids after 0, 10, 20 and 30s low-energy plasma treatment. **D**. A low-magnification micrograph of RGO membrane imaged under defocus condition after irradiation of smaller regions at a total dose of ~200 e^−^ /Å^2^. **E**. A low-magnification micrograph of GO membrane imaged under defocus condition after irradiation of smaller regions (circled by dotted yellow lines) at a total dose of ~200 e^−^/Å^2^. **F**. A micrograph of the same area in **E** taken at overfocus condition. Electron beam-induced footprints are visible in **F** and **E**.

We analyzed the beam-induced effect of RGO and GO. After electron irradiation with an accumulated dose of ~200 e^−^/Å^2^ at room temperature, the RGO membrane had no visible beam-induced footprints (Figure 2D), probably due to its improved electrical conductivity. In contrast, there were obvious beam-induced image marks left on the GO membrane after being irradiated with the same dose, exhibiting as “white disks” in de-focused (Figure 2E) and “black disks” in over-focused micrographs (Figure 2F). These radiation-induced disks may be caused by charging (Brink et al., 1998; Danev et al., 2014; Hettler et al., 2018) or mass loss effects (Choppin et al., 2013; Jiang and Spence, 2009). On the electric-insulating GO membranes, charge could be built up by the accumulated electrons, generating a phase contrast that can be observed in out-of-focus micrographs (Figure 2E-F). On the other hand, small molecules and functional groups sandwiched among GO layers are dose-sensitive and susceptible to electron-beam bombardment (Moon et al., 2010; Zhang et al., 2009), thereby causing decomposition at the exposed regions.

### RGO for Cryo-EM reconstructions

We applied RGO grids to prepare cryo-specimens of 20S proteasome and ribosome, and collected datasets using the same parameters on a Tecnai Arctica microscope (200kV) with Falcon II camera (Figure 3). The particles of both biomolecules were distributed as monodispersed and high-contrast particles on RGO supporting membranes (Figure 3A and 3D). Radiation damage induced by the electron beam is one of the key concerns to be considered in cryo-EM imaging. High-resolution content, i.e., the chemical bonds in biomolecules, is hypothesized to be destroyed first by electron dose during EM imaging. B factor is proposed in cryo-EM field to model the dose and radiation damage on high-resolution information of biomolecules. Here, we ploted B factors of proteasome and ribosome supported by RGO grids with accumulated electron dose (Figure 3B and 3E). The B factors of the first 1-5 e^−^/Å^2^ dose were relatively higher than those of the following 5-10 e^−^/Å^2^, which was mainly resulted from the intial beam-induced motion (Scheres, 2016). We calculated the decay rates of B factor after the first 10 e^−^/Å^2^ of proteasome and ribosome, both of which were approximately 8 Å^2^/(e^−^/Å^2^). Notably, the B-factor decay rates were close to the radiation damage measurements of protein 2D crystals (Peet et al., 2019).

**Figure 3.**
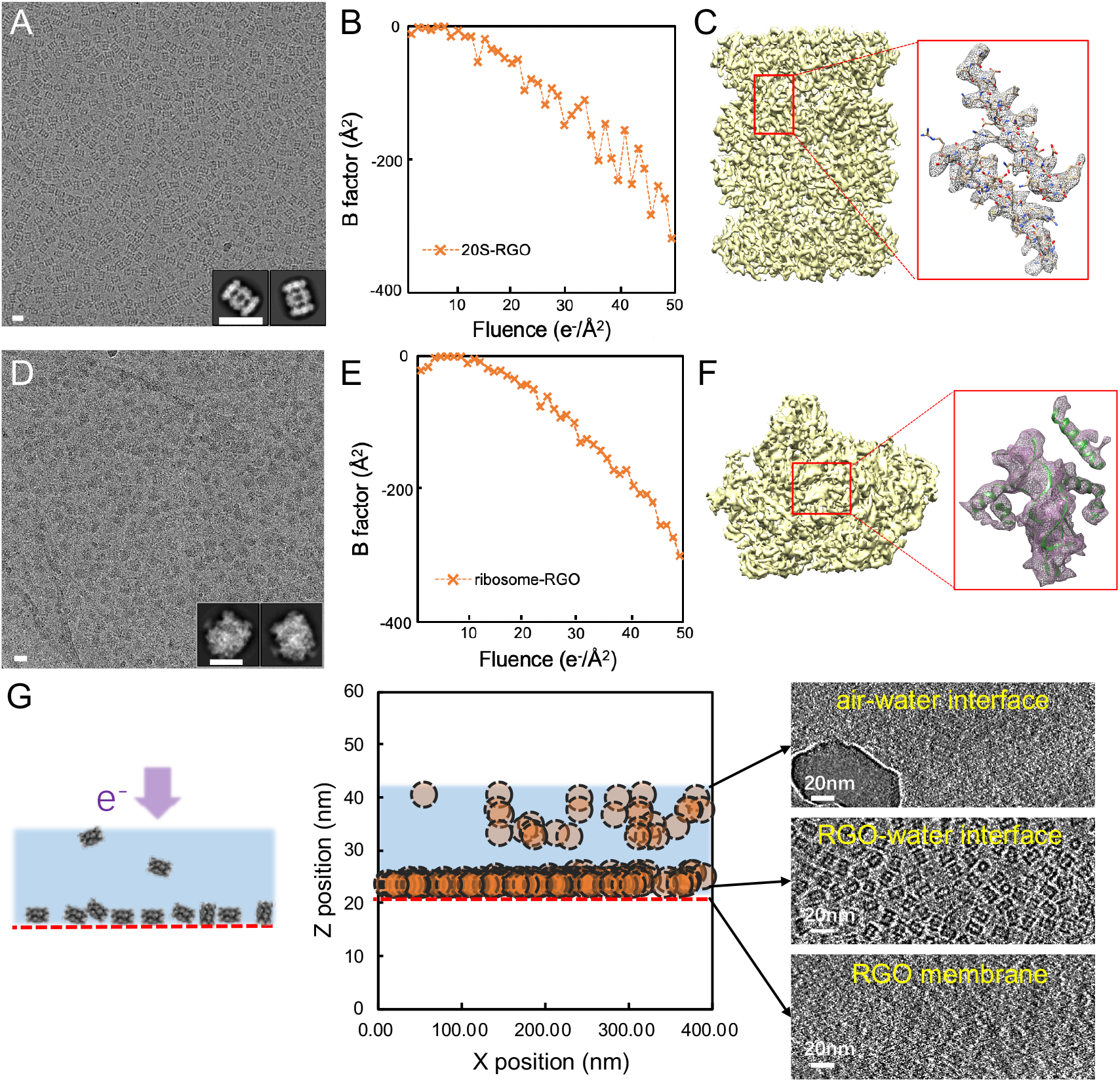
RGO for cryo-EM reconstructions. **A**. A representative micrograph of 20S proteasome on RGO membrane. The inset was the two-dimensional classification results. Scale bar represented 20 nm. **B**. The B factor of the reconstruction of 20S proteasome on RGO was plotted with the accumulated dose. **C**. The reconstructed cryo-EM map of 20S proteasome. The density detail with the corresponding atomic model (PDB: 3J9I (Li et al., 2013)) docked was listed aside in the red box. Some side chains could be clearly recognized. **D**. A representative micrograph of ribosome on RGO membrane. Scale bar represented 20 nm. **E**. The B factor of ribosome plotted with the accumulated dose. **F**. The reconstructed cryo-EM map of ribosome. The density detail with the corresponding atomic model (PDB: 5H4P (Ma et al., 2017)) docked was listed aside in the red box, consistent with the reported resolution. **G**. Cryo-electron tomography of 20S proteasome specimen supported by RGO membrane. Left: the schematic diagram of 20S proteasome particles on RGO membrane, indicated by red dotted line. Middle: the distribution of particles position in ice. The dotted circles indicated individual particles, and the red dotted line indicated RGO membrane. Right: three selected cross sections of the cryo-specimen ET reconstruction. From up to bottom were air-water interface, RGO-water interface and RGO membrane sections, respectively. 20S proteasome particles were mainly distributed on the RGO-water interface.

We also prepared cryo-EM specimens of 20S proteasome and ribosome using GO grids. Interestingly, after processing the three-dimensional reconstruction, the Euler angle distribution of particles on RGO was slightly different from that on GO. There were certain portion of 20S proteasome particles with top view (circle shape) on GO membrane (Figure S5), which were totally absent on RGO (Figure 3A). Since 20S proteasome is highly symmetrical, these side-view projections on RGO were enough to reconstruct its structure. We finally obtained the 20S proteasome reconstruction at 4.4 Å resolution on RGO and 4.7 Å resolution on GO, using the same number of particle images. For ribosome sample of no symmetry, distribution of particle orientations was more balanced on RGO membrane (Figure S6a and c), thus enabling us to get a reconstruction of the ribosome with more reliable structural features than that determined by using particles on GO membranes (Figure 3C, right and Figure S6). We further plotted the directional FSC (Figure S7) (Tan et al., 2017) and calculated the cryoEF value (Naydenova and Russo, 2017), which was 0.28 on GO grid, while 0.42 on RGO grid. Both the directional FSC plots and cryoEF values indicated that ribosome particles on RGO adopted richer orientations. Such variation of particle-orientation performance was probably due to the distinct interacting favor of the biomolecular local surface with the functional groups on supporting membranes whose composition and distribution are different between GO and RGO surfaces. The overall resolution of ribosome reconstruction was reported as 6.1 Å on RGO, which was 8.0 Å on GO (Figure S8 and S9).

Furthermore, we evaluated the protein particle’s distribution in the vitreous ice on RGO-supported specimen using cryo-electron tomography (cryo-ET) technique and found that the majority of particles were absorbed onto RGO surface, thus avoiding damage by the air-water interface (Figure 3G). The ice thickness of the tomogram reconstruction was estimated to be ~20 nm, indicative of little extra background noise generated by the ambient ice.

### Cryo-EM analysis of protein molecules of small molecular weight on RGO grid

From the above analysis, we reasoned that the RGO grid should benefit more to the cryo-EM analysis of smaller macromolecules. We therefore further tested the application of RGO grids on cryo-EM imaging of small biomolecules, such as a 60-kDa protein glycosyltransferase (Figure S10a) and a 50-kDa protein Rv2466c (Figure S10b), both demonstrating particle images of monodisperse and good contrast. Notably, using the RGO grids, Bai et al reconstructed the cryo-EM structure of DEAH-box ATPase/helicase Prp2 (~100 kDa) at a better-than-3-Å resolution, revealing the role of Prp2 in RNA translocation and spliceosome remodeling (Bai et al., 2020).

COVID-19 virus was identified as a novel pathogenic coronavirus emerging to be an enormous threat to the global public health. Receptor binding domain (RBD) of its spike protein exhibited high binding affinity to the ACE2 receptor on human cells (Zhou et al., 2020). We applied RGO grids to prepare the cryo-specimen of RBD-ACE2 complex (95 kDa) and can unambiguously recognize monodispersed particles with good density and contrast on RGO-film-covered area where the graphene diffraction spots were clearly revealed in the Fourier transform (Figure S11a). The contrast of protein particles absorbed on RGO film was further improved when utilizing volta phase plate in cryo-EM (Figure 4A). In contrast, in the area where RGO film was broken (i.e., without RGO membrane support), the RBD-ACE2 particles density was significantly decreased and many extra noisy spots of smaller size compared with target particles were observed in the background (Figure 4B and Figure S13b), probably composed of denatured proteins or dissociated components due to the air-water interface interaction (Figure 3G). From these particle images, we were able to obtain reconstruction of the RBD-ACE2 complex at 2.8 Å resolution and build the atomic model of RBD-ACE2 complex (Figure 4C-F). The interaction interface of ACE2 and RBD of spike protein was clearly recognized in the cryo-EM density (Figure 4E). Interestingly, we found that the cryo-EM structure was more compact than the previously reported structure of the same complex solved by X-ray crystallography (Figure 4G-I) (Lan et al., 2020). There are two possibilities that might lead to such difference. One possibility is that a distinct conformation is favored in the crystal packing of RBD-ACE2 complex during crystallization. While in cryo-EM reconstruction, particles are supposed to be in solution, thus adopting a different conformation. The other possibility is that the adsorption of RBD-ACE2 particles to RGO surface might trap the complex at a specific state. Taken together, the RGO grid was suitable for better protein preservation and cryo-EM analysis of some small biomolecules.

**Figure 4.**
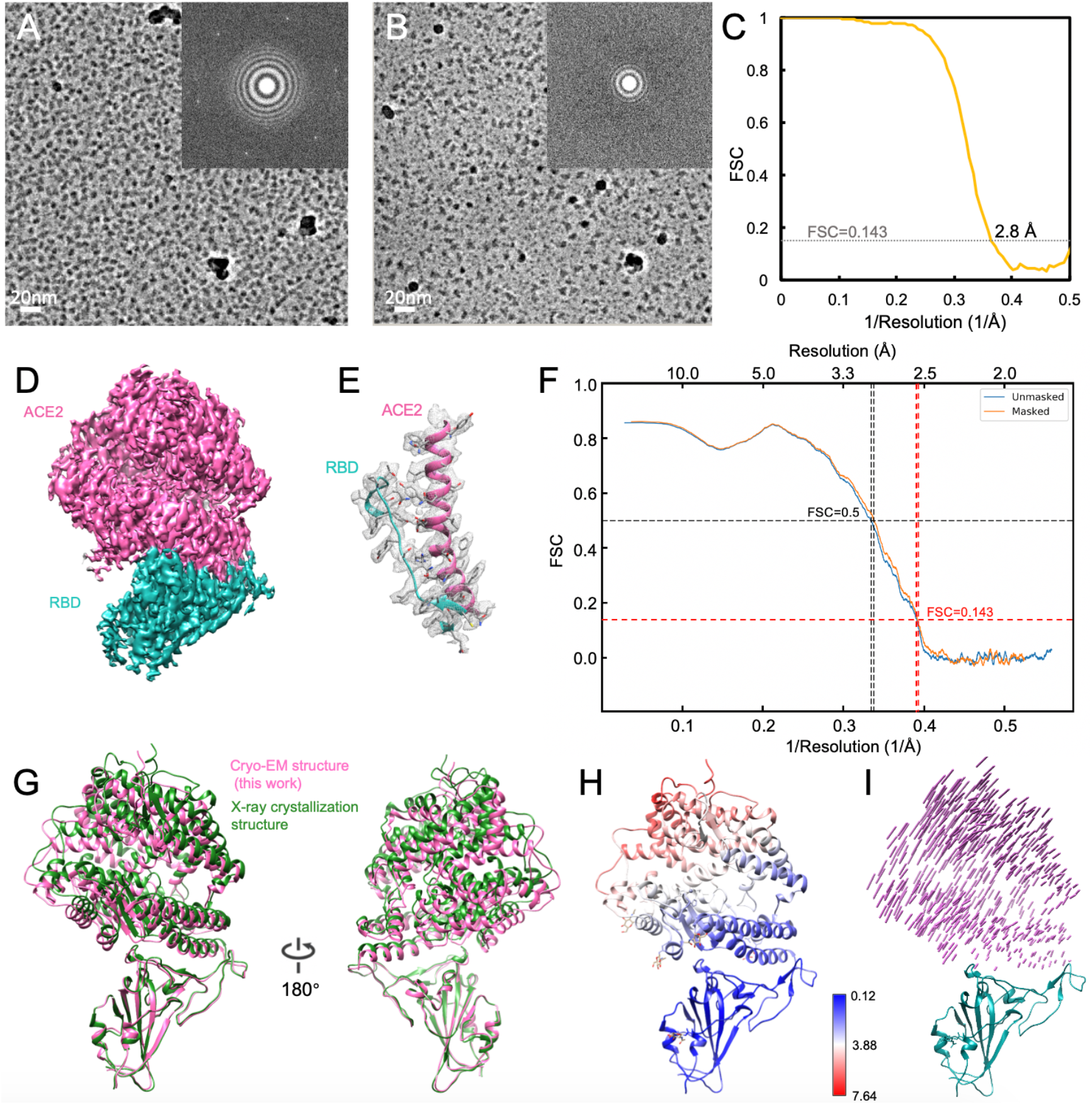
Cryo-EM reconstruction of ACE2-RBD complex supported by RGO membrane. **A.** A representative micrograph of RBD-ACE2 complex in the RGO supporting area. The inset was the corresponding FFT image, where six first-order diffraction spots of graphene were sharply displayed. **B**. A representative micrograph of RBD-ACE2 complex in the RGO-broken area. **C**. The FSC curve of ACE2-RBD complex reconstruction. The dotted gray line indicated FSC=0.143, which was used for estimating the resolution. **D**. The cryo-EM density of ACE2-RBD complex. The pink region was ACE2, while the cyan was RBD. **E**. The ACE2-RBD interaction interface with their atomic models docked. **F**. The ACE2-RBD model-map FSC curve. **G**. Comparison between the cryo-EM structure solved in this work (colored in pink) with the reported X-ray crystal structure (PDB: 6M0J, in green). **H**. RMSD (Å) between the cryo-EM structure and the X-ray crystal structure. **I**. 3D vector map between the cryo-EM structure and the X-ray crystal structure, using RBD as the alignment reference. The vector lengths indicated the displacement scale across these two structures.

## Conclusion

Graphene membrane has long been explored as the supporting film in electron microscopy (Wilson et al., 2009). The sp^2^ carbon-atom crystalline lattice enables graphene ultrahigh electrical/thermal conductivity. The single-atomic thickness much smaller than the mean free path of electron presents negligible background noise. Yet its practical application is largely limited by the difficulty of reproducible high-quality graphene synthesis and free-of-contamination transfer onto EM grids. Here we described a robust method to fabricate single- or few-layer reduced graphene oxide membrane coated EM grids in large scale. We demonstrated that RGO contained decreased interlayer space and fewer functional groups, compared with GO. Importantly, we found that RGO can recover the electrical conductivity to a certain extent and the capability of reducing charging effects, thus improving the image stability. RGO was able to keep particles on its surface and more friendly than the air-water interface. We finally applied RGO grids for cryo-EM specimen preparation of sub-100 kDa protein samples to reconstruct the small molecules at near atomic resolution. What needs to be pointed out here is that adsorption of particles onto the surface of graphene supporting film often resulted in preferred orientation or other unfavorable consequences that still limited the resolution that could be achieved. Better reconstruction results might be obtained by using RGO with multiple bioactive functionalization.

## Materials and Methods

### Preparation of graphene oxide solution

The graphene oxide solution was purchased from Sigma-Aldrich company (Lot #777676) or prepared by improved Hummer’s method (Marcano et al., 2010). Briefly, 9g KMnO_4_ and 1.5 g graphite flakes were added and mixed into 200 mL acid solvent (H_2_SO_4_/H_3_PO_4_ with a volume ratio of 9:1). The mixed solution was then stirred for 12 hours at 50°C. Afterwards, the solution was cooled down in ice bath and was added with 200 mL ice-cold water containing 3 mL H_2_O_2_. After 2-hour standing on ice, the mixture was centrifuged at 5000 g for 20 min, and the precipitate was carefully collected and washed, and finally resuspended in distilled water to make graphene oxide (GO) solution.

### Fabrication of GO grids

Graphene oxide membrane was transferred onto Quantifoil gold EM grids by a similar procedure described by Palovcak et al. (Palovcak et al., 2018). Briefly, the GO solution was first mixed into the dispersant (methanol/water with a volume ratio of 5:1) and sonicated for 10 min. The solution was next centrifuged at 4000 g for 10 min and the pellets were resuspended in the dispersant solution, followed by a 2-min sonication. Then the GO solution was centrifuged at 500 g for 1 min, and the supernatant was carefully collected for coating EM grids. When fabricating GO EM grids, we firstly submerge a steel mesh stand covered by a piece of filter paper into a water-filled container with outlets at its bottom (Supplementary Figure 1B). The Quantifoil gold EM grids were mildly glow-discharged in advance and put onto the steel stand. GO solution was gently pipetted onto the water bath surface to form GO film and the water was slowly drained to lay the GO film onto the grid surface. Finally, the EM grids supported by the filter paper were carefully taken out from the container and air-dried at a 60°C baker.

### Preparation of RGO grids

The as-fabricated GO grids were placed in a quartz boat and put into a clean tube furnace at room temperature. A flow of 100 sccm H_2_ and 100 sccm Ar was introduced to drain the air out of the furnace. The GO grids were then heated to 300°C at a rate of 2°C/min, under the atmosphere of 100 sccm H_2_ and 100 sccm Ar. Subsequently, the grids were reduced at 300°C for 1 hour under the same atmosphere. After reduction, the grids were naturally cooled down to room temperature and ready for use.

### Characterization

The morphologies of the GO and RGO membranes were characterized using AFM (Bruker dimension icon, scansyst mode, scansyst air tip). The composition characterizations of GO and RGO membranes were conducted with XPS (Kratos Analytical AXIS-Ultra with monochromatic Al Kα X-ray). The four-probe resistance measuring meter (CDE ResMap 178) was used to measure the sheet resistances of GO and RGO membranes on mica substrates. The graphene layer number was determined by the diffraction pattern under TEM. The first-order diffraction pattern of single-layer graphene contained six spots, the density of which were similar with that of the second-order diffraction spots. The water contact angle of RGO and GO grids after plasma treatment was measured by the optical contact angle measuring device (Dataphysics company).

### The Sheet resistance measurement

To measure the sheet resistance, four-point probes were equally spaced and arranged in a line on the target material. By measuring the current (I) through the outside two probes and the voltage (V) across the inside two probes, the average sheet resistance (R_s_) of the conducting film can be acquired. When the thickness of the conducting film is much less than the spacing of probes, and the width of film is apparently larger than the spacing distance, the average sheet resistance can be given by R_s_=4.53V/I.

### Cryo-EM specimen preparation

*Thermoplasma acidophilum* 20S proteasome was recombinantly expressed in *Escherichia coli* cells and purified as described previously (Li et al., 2013). Yeast ribosomes were purified from yeast cells following previously published protocols (Ma et al., 2017). RBD-ACE2 complex was prepared according to methods utilized in (Lan et al., 2020). ~3μL solution containing purified biomolecules was pipetted onto freshly glow discharged RGO grids, and then transferred into an FEI Vitrobot. For glow discharging, we used low-energy plasma to treat the RGO grids for 15s in a Harrick PDC-32G plasma cleaner. The humidity of Vitrobot chamber was kept as 100% and the temperature as 8°C. The grids were then blotted 2s with −2 force, immediately followed by plunge-freezing into liquid ethane cooled at liquid nitrogen temperature. After that, the grids were quickly transferred into liquid nitrogen for storage.

### Cryo-EM data collection and analysis

The single-particle cryo-EM datasets of 20S proteasome and ribosome were collected on an FEI Tecnai Arctica (200 kV), equipped with an Falcon II detector. The RBD-ACE2 dataset was collected on an FEI Titan Krios (300 kV), equipped with a Gatan K3 summit detector and Volta phase plate, using AutoEMation software (Fan et al., 2017; Lei and Frank, 2005). All micrographs were dose-dependently fractionated into 32 frames, with an accumulated dose of 50 e^−^/Å^2^. The individual frames were firstly motion-corrected by MotionCor2 algorithm (Zheng et al., 2017) and the resulted micrographs were imported into Relion3.0 (Zivanov et al., 2018) for further processing. CTF values were calculated by CTFFIND4 package (Rohou and Grigorieff, 2015). Biomolecular particles were autopicked and iteratively 2D classified in Relion3.0. Particles grouped in good classes exhibiting fine structural details were selected for further 3D classification and refinement. We used Fourier Shell Correction (FSC) 0.143 cutoff criteria to estimate the resolution in the final postprocessing step in Relion3.0. Finally, we used 10,000 and 8,453 particles for the reconstructions of 20S proteasome and ribosome. The resolution of 20S proteasome and ribosome on RGO grids was 4.5 Å and 6.1 Å, respectively. For RBD-ACE2 complex, we finally used 110,122 particles and got a reconstruction at 2.8 Å resolution. All figures related to cryo-EM structures were created in UCSF Chimera (Pettersen et al., 2004). The directional FSC curves were generated in remote 3DFSC processing server (https://3dfsc.salk.edu) (Tan et al., 2017).

### Cryo-electron tomography analysis

Cryo-ET micrographs series were obtained on an FEI Titan Krios microscope (300 kV), equipped with a Gatan K2 camera. We used SerialEM (Mastronarde, 2005) to collect tilt series, and the specimen was tilted from +51° to −51° with an acquired step of 3°. The total dose was 100 e^−^/Å^2^ with a dose rate of ~3 e^−^/Å^2^/s, and the calibrated pixel size was 1.77 Å. The micrographs series was imported into IMOD (Kremer et al., 1996) for reconstruction, where the position of protein particles was manually identified as previously (Liu et al., 2019).

## Acknowledgements

We thank Dr. Ning Gao, Dr. Xueming Li for kindly providing ribosome and 20S proteasome samples. We are grateful to Dr. Jianlin Lei, Dr. Lingpeng Cheng, Dr. Tao Yang, Dr. Xiaomin Li, Dr. Fan Yang, Danyang Li, Xiaofeng Hu, Jie Wen, Yakun Wang, and Anbao Jia at the Cryo-EM and High-Performance Computation platforms of Tsinghua University Branch of the National Protein Science Facility, for the technical support in cryo-EM data collection and analysis. This work is financially supported by the Ministry of Science and Technology of China (2016YFA0501100) and National Natural Science Foundation of China (31825009) to H.-W.W., and the National Natural Science Foundation of China (21525310) and the National Basic Research Program of China (2014CB932500 and 2016YFA0200101) to H.-L.P..

## Contributions

Y.C., H.-L.P., and H.-W.W. conceived the project. N.L., L.Z., J.X., and J.Z. prepared RGO grids and conduted the physicochemical charateriaztion. N.L., J.X., J.W., C.H., J.L., and H.C. prepared the cryo-EM specimens and processed the single-particle reconstruction. N.L., X.Z., K.X., and Z.Y. performed the cryo-ET analysis. N.L., L.Z., H.-L.P, Y.C., and H.-W.W. wroted the manuscript.

## Data and material availability

The cryo-EM map of SARS-CoV2 RBD-ACE2 complex has been deposited in the EMDB under accession number EMD-30816, and the coordinate in the PDB with PDB ID 7DQA.

## Compliance with ethics guidelinnes

### Human and animal rights and informed consent

The article does not contain any studies with human or animal subjects performed by the any of the authors.

## Reference

Armstrong, M., Han, B.G., Gomez, S., Turner, J., Fletcher, D.A., and Glaeser, R.M. (2020). Microscale Fluid Behavior during Cryo-EM Sample Blotting. Biophys J 118, 708–719.

Bai, R., Wan, R., Yan, C., Jia, Q., Lei, J., and Shi, Y. (2020). Mechanism of spliceosome remodeling by the ATPase/helicase Prp2 and its coactivator Spp2. Science.

Balandin, A.A., Ghosh, S., Bao, W.Z., Calizo, I., Teweldebrhan, D., Miao, F., and Lau, C.N. (2008). Superior thermal conductivity of single-layer graphene. Nano Lett 8, 902–907.

Benjamin, C.J., Wright, K.J., Bolton, S.C., Hyun, S.H., Krynski, K., Grover, M., Yu, G.M., Guo, F., Kinzer-Ursem, T.L., Jiang, W., et al. (2016). Selective Capture of Histidine-tagged Proteins from Cell Lysates Using TEM grids Modified with NTA-Graphene Oxide. Sci Rep-Uk 6.

Brink, J., Sherman, M.B., Berriman, J., and Chiu, W. (1998). Evaluation of charging on macromolecules in electron cryomicroscopy. Ultramicroscopy 72, 41–52.

Buchsteiner, A., Lerf, A., and Pieper, J. (2006). Water dynamics in graphite oxide investigated with neutron scattering. J Phys Chem B 110, 22328–22338.

Chen, J.H., Jang, C., Adam, S., Fuhrer, M.S., Williams, E.D., and Ishigami, M. (2008). Charged-impurity scattering in graphene. Nat Phys 4, 377–381.

Cheng, Y. (2015). Single-Particle Cryo-EM at Crystallographic Resolution. Cell 161, 450–457.

Cheng, Y.F. (2018). Single-particle cryo-EM-How did it get here and where will it go. Science 361, 876–+.

Choppin, G., Liljenzin, J.-O., Rydberg, J., and Ekberg, C. (2013). Absorption of Nuclear Radiation. In Radiochemistry and Nuclear Chemistry, pp. 163–208.

D’Imprima, E., Floris, D., Joppe, M., Sanchez, R., Grininger, M., and Kuhlbrandt, W. (2019). Protein denaturation at the air-water interface and how to prevent it. Elife 8.

Danev, R., Buijsse, B., Khoshouei, M., Plitzko, J.M., and Baumeister, W. (2014). Volta potential phase plate for in-focus phase contrast transmission electron microscopy. P Natl Acad Sci USA 111, 15635–15640.

Dubochet, J., Lepault, J., Freeman, R., Berriman, J.A., and Homo, J.C. (1982). Electron-Microscopy of Frozen Water and Aqueous-Solutions. J Microsc-Oxford 128, 219–237.

Egerton, R.F., Li, P., and Malac, M. (2004). Radiation damage in the TEM and SEM. Micron 35, 399–409.

Fan, X., Zhao, L., Liu, C., Zhang, J.C., Fan, K., Yan, X., Peng, H.L., Lei, J., and Wang, H.W. (2017). Near-Atomic Resolution Structure Determination in Over-Focus with Volta Phase Plate by Cs-Corrected Cryo-EM. Structure 25, 1623–1630 e1623.

Geim, A.K., and Novoselov, K.S. (2007). The rise of graphene. Nat Mater 6, 183–191.

Glaeser, R.M. (2016). Specimen Behavior in the Electron Beam. Methods Enzymol 579, 19–50.

Glaeser, R.M. (2018). Proteins, Interfaces, and Cryo-Em Grids. Curr Opin Colloid Interface Sci 34, 1–8.

Grassucci, R.A., Taylor, D.J., and Frank, J. (2007). Preparation of macromolecular complexes for cryo-electron microscopy. Nature Protocols 2, 3239–3246.

Han, Y., Fan, X., Wang, H., Zhao, F., Tully, C.G., Kong, J., Yao, N., and Yan, N. (2020). High-yield monolayer graphene grids for near-atomic resolution cryoelectron microscopy. Proc Natl Acad Sci U S A 117, 1009–1014.

Hettler, S., Kano, E., Dries, M., Gerthsen, D., Pfaffmann, L., Bruns, M., Beleggia, M., and Malac, M. (2018). Charging of carbon thin films in scanning and phase-plate transmission electron microscopy. Ultramicroscopy 184, 252–266.

Jiang, N., and Spence, J.C.H. (2009). Radiation damage in zircon by high-energy electron beams. J Appl Phys 105.

Jung, I., Dikin, D.A., Piner, R.D., and Ruoff, R.S. (2008). Tunable Electrical Conductivity of Individual Graphene Oxide Sheets Reduced at “Low” Temperatures. Nano Lett 8, 4283–4287.

Kremer, J.R., Mastronarde, D.N., and McIntosh, J.R. (1996). Computer visualization of three-dimensional image data using IMOD. Journal of Structural Biology 116, 71–76.

Lan, J., Ge, J., Yu, J., Shan, S., Zhou, H., Fan, S., Zhang, Q., Shi, X., Wang, Q., Zhang, L., et al. (2020). Structure of the SARS-CoV-2 spike receptor-binding domain bound to the ACE2 receptor. Nature.

Lee, C., Wei, X.D., Kysar, J.W., and Hone, J. (2008). Measurement of the elastic properties and intrinsic strength of monolayer graphene. Science 321, 385–388.

Lei, J., and Frank, J. (2005). Automated acquisition of cryo-electron micrographs for single particle reconstruction on an FEI Tecnai electron microscope. J Struct Biol 150, 69–80.

Li, X., Mooney, P., Zheng, S., Booth, C.R., Braunfeld, M.B., Gubbens, S., Agard, D.A., and Cheng, Y. (2013). Electron counting and beam-induced motion correction enable near-atomic-resolution single-particle cryo-EM. Nat Methods 10, 584–590.

Lian, B., De Luca, S., You, Y., Alwarappan, S., Yoshimura, M., Sahajwalla, V., Smith, S.C., Leslie, G., and Joshi, R.K. (2018). Extraordinary water adsorption characteristics of graphene oxide. Chem Sci 9, 5106–5111.

Lin, L., Sun, L.Z., Zhang, J.C., Sun, J.Y., Koh, A.L., Peng, H.L., and Liu, Z.F. (2016). Rapid Growth of Large Single-Crystalline Graphene via Second Passivation and Multistage Carbon Supply. Advanced Materials 28, 4671–4677.

Liu, N., Zhang, J., Chen, Y., Liu, C., Zhang, X., Xu, K., Wen, J., Luo, Z., Chen, S., Gao, P., et al. (2019). Bioactive Functionalized Monolayer Graphene for High-Resolution Cryo-Electron Microscopy. J Am Chem Soc 141, 4016–4025.

Ma, C.Y., Wu, S., Li, N.N., Chen, Y., Yan, K.G., Li, Z.F., Zheng, L.Q., Lei, J.L., Woolford, J.L., and Gao, N. (2017). Structural snapshot of cytoplasmic pre-60S ribosomal particles bound by Nmd3, Lsg1, Tif6 and Reh1. Nature Structural & Molecular Biology 24, 214–+.

Marcano, D.C., Kosynkin, D.V., Berlin, J.M., Sinitskii, A., Sun, Z., Slesarev, A., Alemany, L.B., Lu, W., and Tour, J.M. (2010). Improved synthesis of graphene oxide. ACS Nano 4, 4806–4814.

Mastronarde, D.N. (2005). Automated electron microscope tomography using robust prediction of specimen movements. Journal of Structural Biology 152, 36–51.

Moon, I.K., Lee, J., Ruoff, R.S., and Lee, H. (2010). Reduced graphene oxide by chemical graphitization. Nat Commun 1, 73.

Nakane, T., Kotecha, A., Sente, A., McMullan, G., Masiulis, S., Brown, P.M.G.E., Grigoras, I.T., Malinauskaite, L., Malinauskas, T., Miehling, J., et al. (2020).

Naydenova, K., Peet, M.J., and Russo, C.J. (2019). Multifunctional graphene supports for electron cryomicroscopy. Proc Natl Acad Sci U S A 116, 11718–11724.

Naydenova, K., and Russo, C.J. (2017). Measuring the effects of particle orientation to improve the efficiency of electron cryomicroscopy. Nat Commun 8, 629.

Noble, A.J., Wei, H., Dandey, V.P., Zhang, Z., Tan, Y.Z., Potter, C.S., and Carragher, B. (2018). Reducing effects of particle adsorption to the air-water interface in cryo-EM. Nat Methods 15, 793–795.

Palovcak, E., Wang, F., Zheng, S.Q., Yu, Z., Li, S., Betegon, M., Bulkley, D., Agard, D.A., and Cheng, Y. (2018). A simple and robust procedure for preparing graphene-oxide cryo-EM grids. J Struct Biol 204, 80–84.

Pantelic, R.S., Meyer, J.C., Kaiser, U., Baumeister, W., and Plitzko, J.M. (2010). Graphene oxide: a substrate for optimizing preparations of frozen-hydrated samples. J Struct Biol 170, 152–156.

Peet, M.J., Henderson, R., and Russo, C.J. (2019). The energy dependence of contrast and damage in electron cryomicroscopy of biological molecules. Ultramicroscopy 203, 125–131.

Pettersen, E.F., Goddard, T.D., Huang, C.C., Couch, G.S., Greenblatt, D.M., Meng, E.C., and Ferrin, T.E. (2004). UCSF chimera - A visualization system for exploratory research and analysis. J Comput Chem 25, 1605–1612.

Qiu, T.F., Luo, B., Liang, M.H., Ning, J., Wang, B., Li, X.L., and Zhi, L.J. (2015). Hydrogen reduced graphene oxide/metal grid hybrid film: towards high performance transparent conductive electrode for flexible electrochromic devices. Carbon 81, 232–238.

Regan, W., Alem, N., Aleman, B., Geng, B.S., Girit, C., Maserati, L., Wang, F., Crommie, M., and Zettl, A. (2010). A direct transfer of layer-area graphene. Appl Phys Lett 96.

Rohou, A., and Grigorieff, N. (2015). CTFFIND4: Fast and accurate defocus estimation from electron micrographs. J Struct Biol 192, 216–221.

Russo, C.J., and Passmore, L.A. (2014a). Controlling protein adsorption on graphene for cryo-EM using low-energy hydrogen plasmas. Nat Methods 11, 649–652.

Russo, C.J., and Passmore, L.A. (2014b). Controlling protein adsorption on graphene for cryo-EM using low-energy hydrogen plasmas. Nature Methods 11, 649–+.

Russo, C.J., and Passmore, L.A. (2014c). Ultrastable gold substrates for electron cryomicroscopy. Science 346, 1377–1380.

Scheres, S.H. (2012). RELION: implementation of a Bayesian approach to cryo-EM structure determination. J Struct Biol 180, 519–530.

Scheres, S.H. (2016). Processing of Structurally Heterogeneous Cryo-EM Data in RELION. Methods Enzymol 579, 125–157.

Tan, Y.Z., Baldwin, P.R., Davis, J.H., Williamson, J.R., Potter, C.S., Carragher, B., and Lyumkis, D. (2017). Addressing preferred specimen orientation in single-particle cryo-EM through tilting. Nat Methods 14, 793–796.

Wang, F., Yu, Z., Betegon, M., Campbell, M.G., Aksel, T., Zhao, J., Li, S., Douglas, S.M., Cheng, Y., and Agard, D.A. (2020). Amino and PEG-amino graphene oxide grids enrich and protect samples for high-resolution single particle cryo-electron microscopy. Journal of Structural Biology 209.

Wang, Y.L., Chen, Y.A., Lacey, S.D., Xu, L.S., Xie, H., Li, T., Danner, V.A., and Hu, L.B. (2018). Reduced graphene oxide film with record-high conductivity and mobility. Mater Today 21, 186–192.

Wilson, N.R., Pandey, P.A., Beanland, R., Young, R.J., Kinloch, I.A., Gong, L., Liu, Z., Suenaga, K., Rourke, J.P., York, S.J., et al. (2009). Graphene Oxide: Structural Analysis and Application as a Highly Transparent Support for Electron Microscopy. Acs Nano 3, 2547–2556.

Wu, S., Armache, J.P., and Cheng, Y. (2016). Single-particle cryo-EM data acquisition by using direct electron detection camera. Microscopy (Oxf) 65, 35–41.

Yip, K.M., Fischer, N., Paknia, E., Chari, A., and Stark, H. (2020).

You, S., Sundqvist, B., and Talyzin, A.V. (2013). Enormous lattice expansion of hummers graphite oxide in alcohols at low temperatures. ACS Nano 7, 1395–1399.

Zhang, J., Lin, L., Sun, L., Huang, Y., Koh, A.L., Dang, W., Yin, J., Wang, M., Tan, C., Li, T., et al. (2017). Clean Transfer of Large Graphene Single Crystals for High-Intactness Suspended Membranes and Liquid Cells. Adv Mater 29.

Zhang, X.Y., Huang, Y., Wang, Y., Ma, Y.F., Liu, Z.F., and Chen, Y.S. (2009). Synthesis and characterization of a graphene-C-60 hybrid material. Carbon 47, 334–337.

Zheng, L., Chen, Y., Li, N., Zhang, J., Liu, N., Liu, J., Dang, W., Deng, B., Li, Y., Gao, X., et al. (2020). Robust ultraclean atomically thin membranes for atomic-resolution electron microscopy. Nat Commun 11, 541.

Zheng, S.Q., Palovcak, E., Armache, J.P., Verba, K.A., Cheng, Y., and Agard, D.A. (2017). MotionCor2: anisotropic correction of beam-induced motion for improved cryo-electron microscopy. Nat Methods 14, 331–332.

Zhou, P., Yang, X.L., Wang, X.G., Hu, B., Zhang, L., Zhang, W., Si, H.R., Zhu, Y., Li, B., Huang, C.L., et al. (2020). A pneumonia outbreak associated with a new coronavirus of probable bat origin. Nature.

Zivanov, J., Nakane, T., Forsberg, B.O., Kimanius, D., Hagen, W.J., Lindahl, E., and Scheres, S.H. (2018). New tools for automated high-resolution cryo-EM structure determination in RELION-3. Elife 7.

