## Supplementary Information for "Reduced graphene oxide membrane as supporting film for high-resolution cryo-EM"

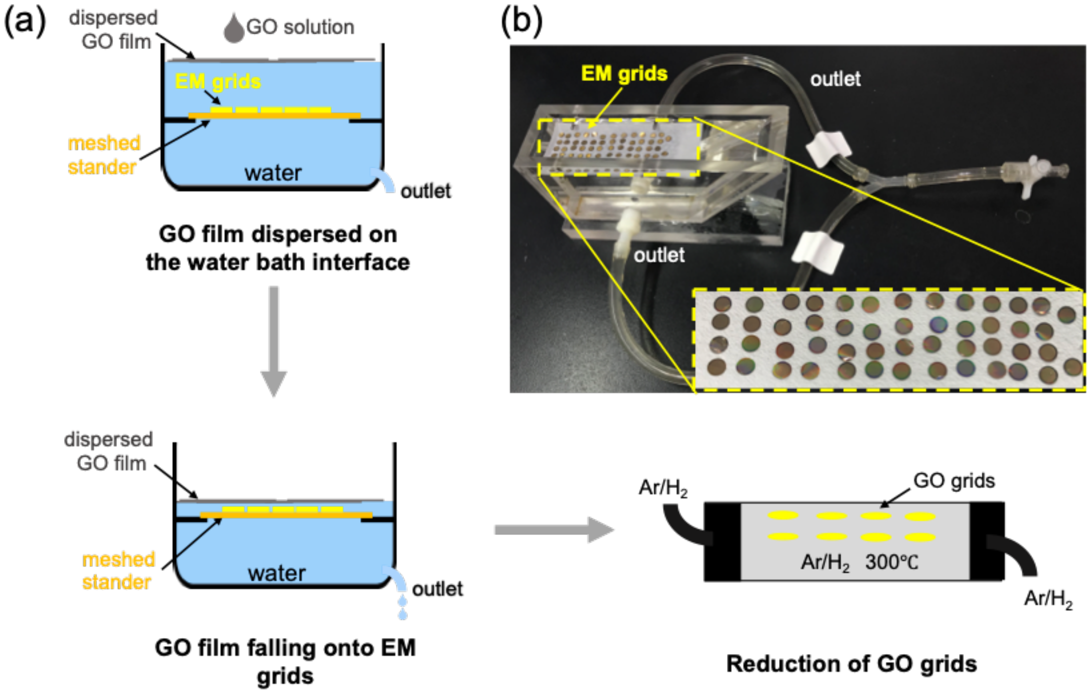


**Figure S1.** Flow diagram of RGO grids fabrication. (a) GO solution was gently pipetted onto the water bath surface, and dispersed to form GO film at the air-water interface. The EM grids had been placed on the meshed steel stander which was immersed into the water bath in advance. With water slowly drained, the GO film would be coated onto EM grids and then air-dried. The GO grids were finally baked in an atmosphere of $H_2$/Ar at 300 °C for 1 hour to produce reduced graphene oxide (RGO) grids. (b) the device used for coating GO film onto EM grids on a large scale.

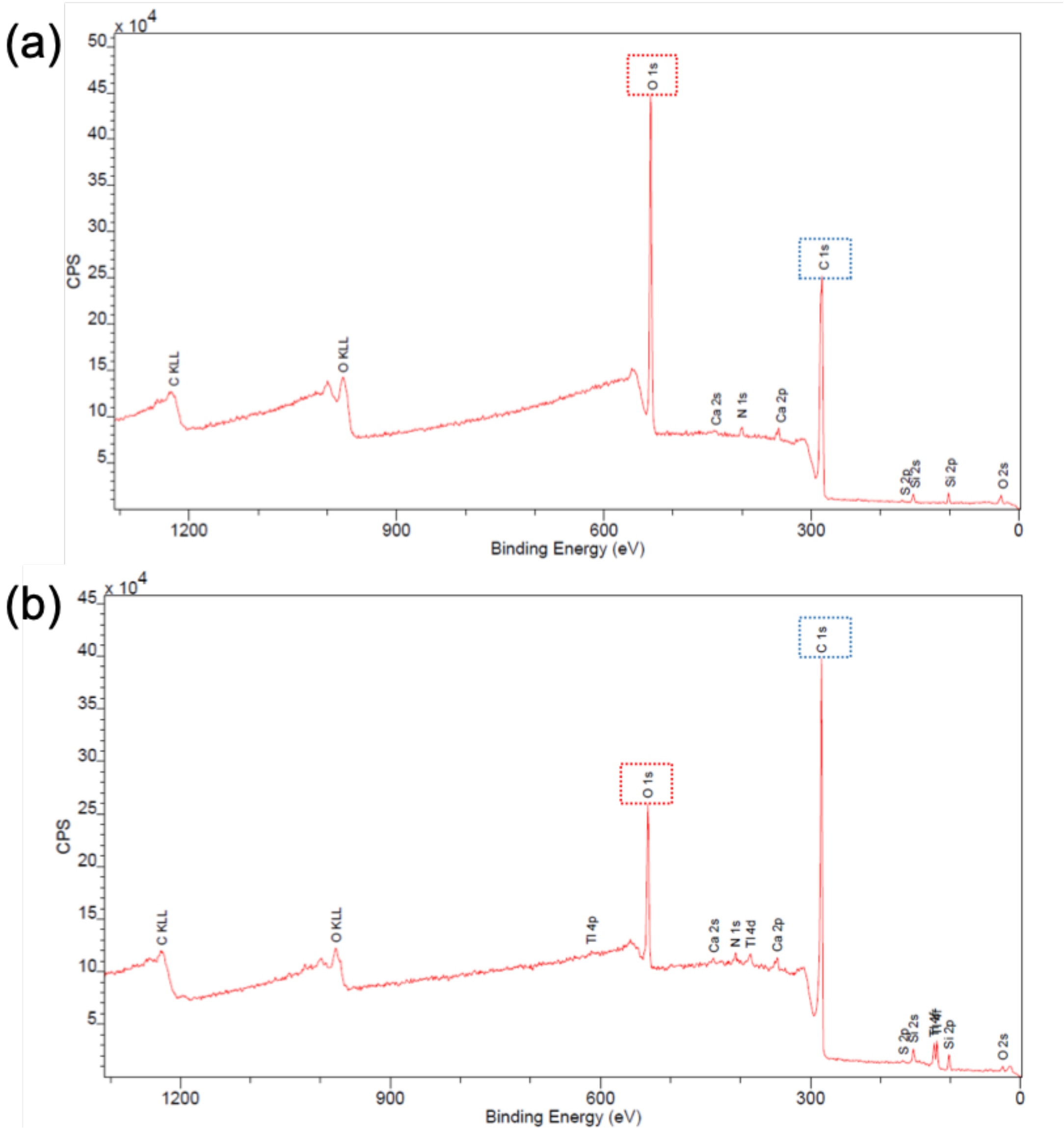


**Figure S2.** XPS survey spectra of GO (a) and RGO (b) grids.

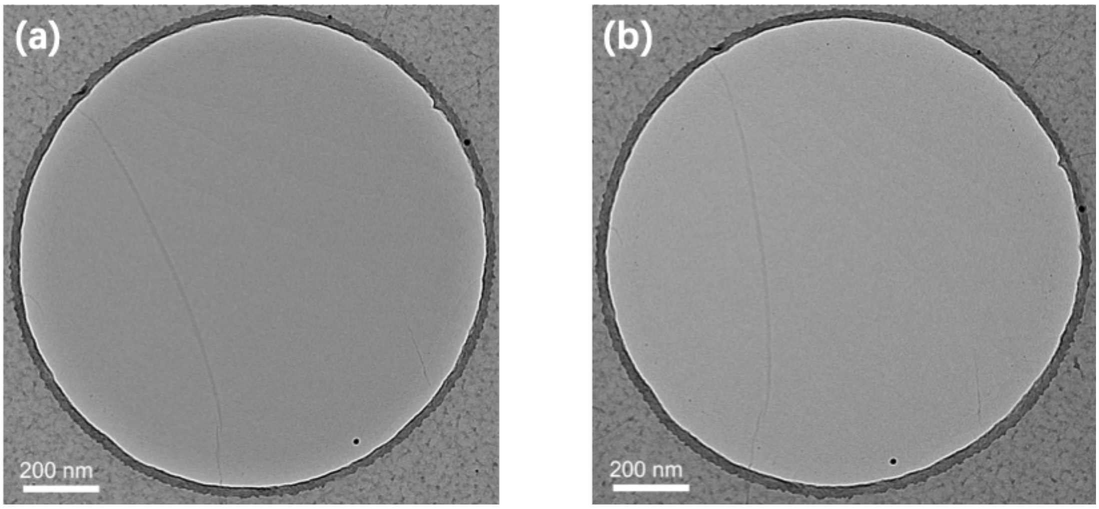


**Figure S3.** TEM micrographs of a GO film coated hole (a) and the same hole after reduction treatment (b).

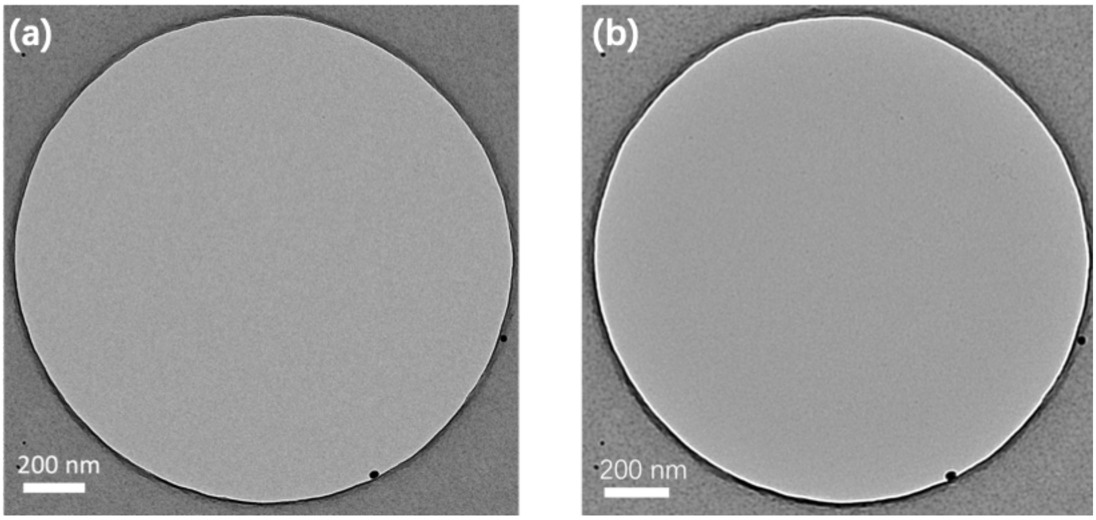


**Figure S4.** TEM micrographs of a RGO film coated hole at a defocus of 3μm (a) and 15μm (b). The RGO film was almost free of contaminations.

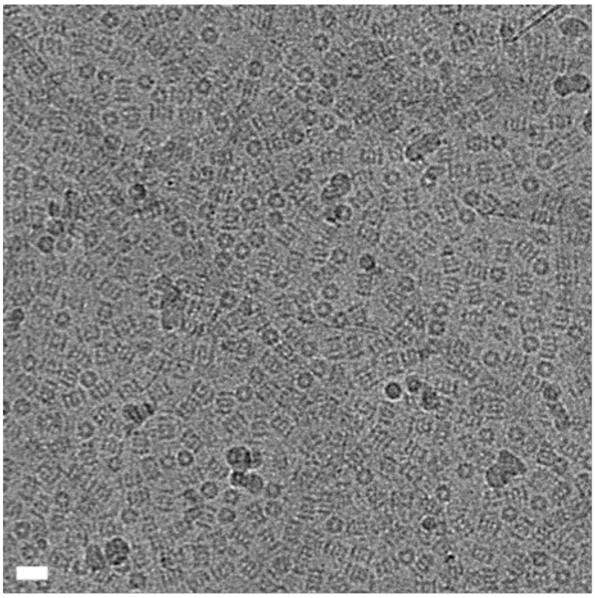

**Figure S5.** A representative cryo-EM micrograph of 20S proteasome on the GO grid. The scale bar represented 20 nm. There were certain portion of 20S proteasome particles with top view (circle shape), which was totally absent in the RGO grid.

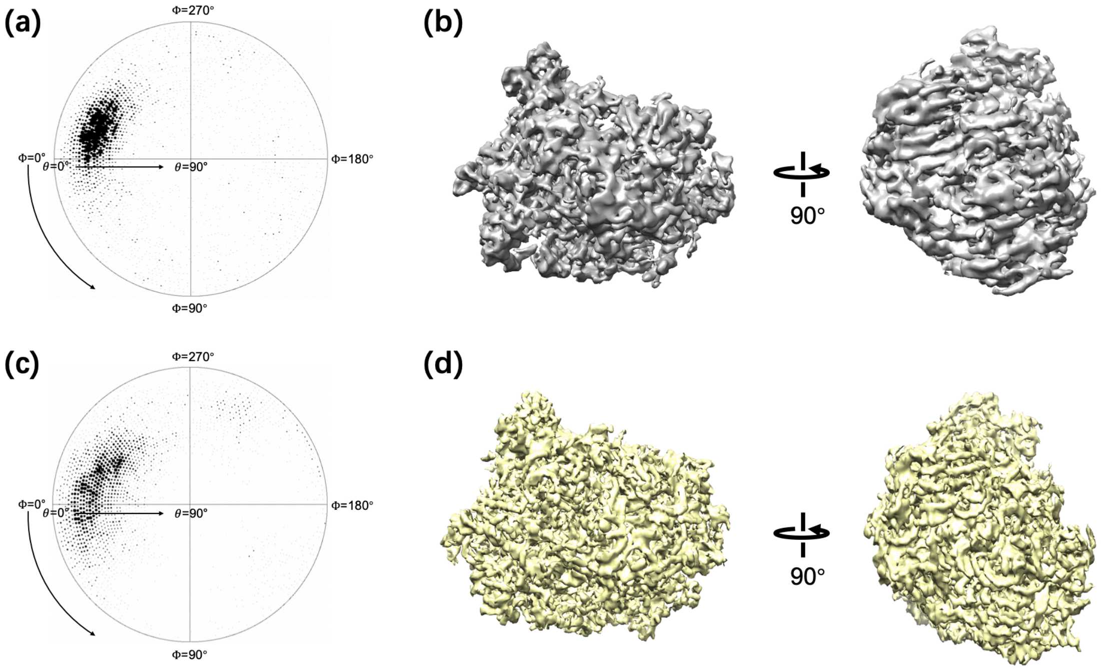


**Figure S6.** (a) the Euler angle distribution of ribosome particles on GO grids. (b) the cryo-EM density of ribosome reconstructed by using particles on GO grids. (c) the Euler angle distribution of ribosome particles on RGO grids. (d) the cryo-EM density of ribosome reconstructed by using particles on RGO grids.

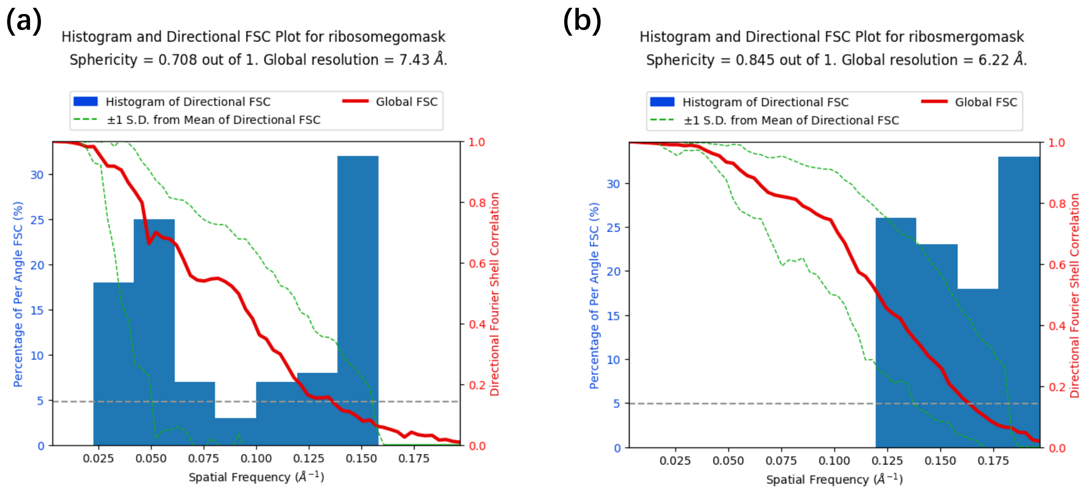


**Figure S7.** The directional FSC curves of the reconstructions of ribosome supported by the GO (a) and RGO (b) grids. The histogram of directional FSC in (a) was more non-uniform than that in (b), indicative of stronger preferred orientation of ribosome particles on the GO grids.

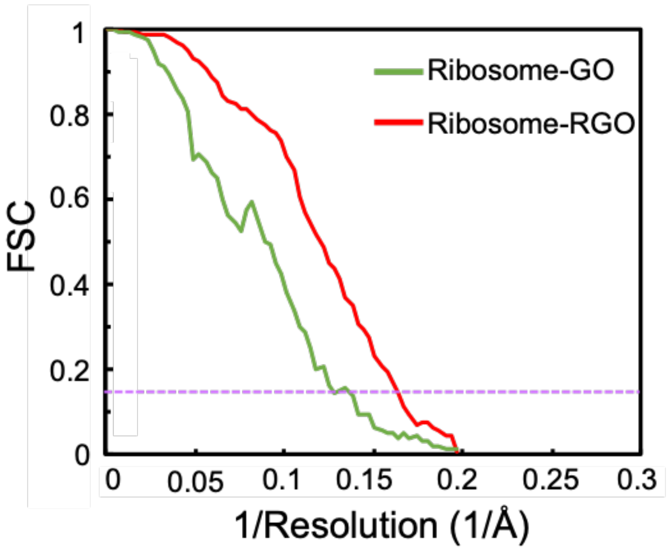


**Figure S8.** FSC curves of ribosome reconstructions on GO and RGO grids. The dotted purple line indicated FSC=0.143.

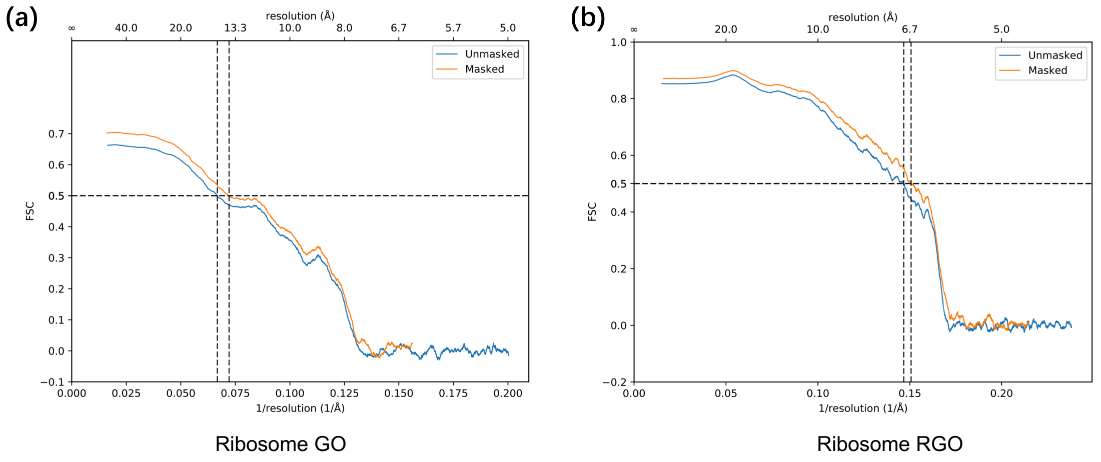


**Figure S9.** The model-to-map FSC curves of ribosome reconstructions on GO (a) and RGO (b).

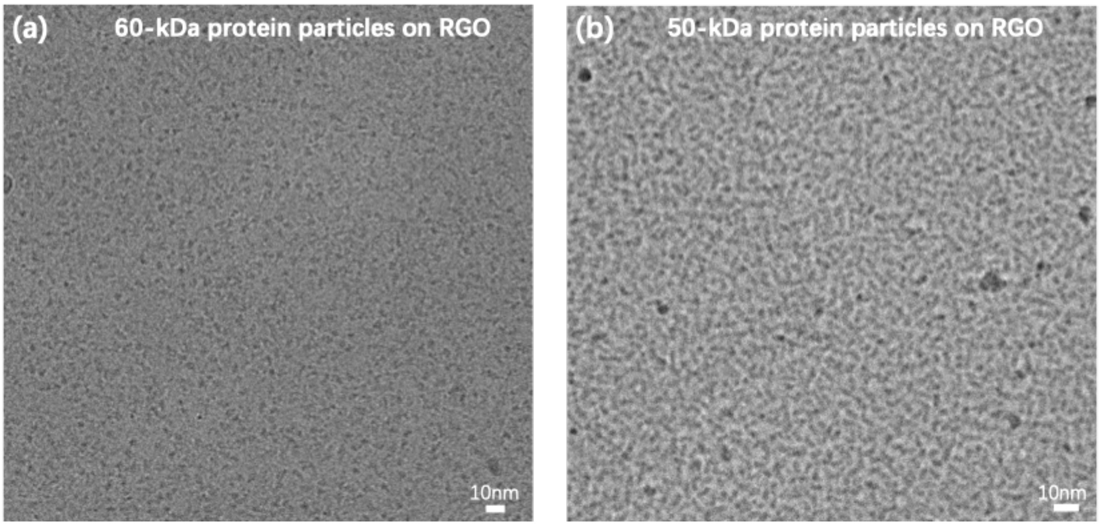


**Figure S10.** Representative cryo-EM micrographs of a 60-kDa protein glycosyltransferase (a) and a 50-kDa protein Rv2466c (b) supported by RGO grids, both exhibiting nice contrast.

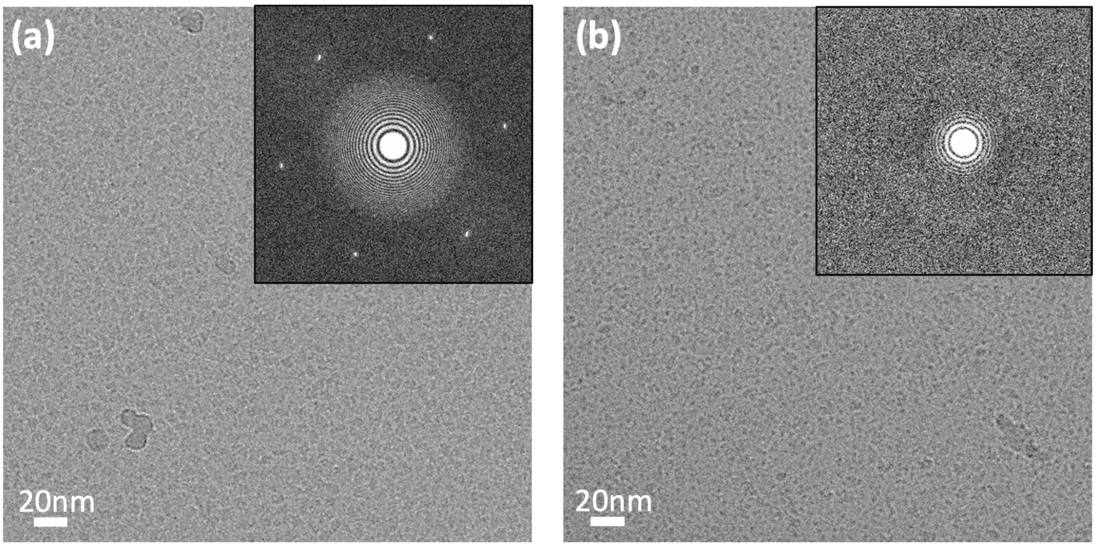


**Figure S11.** Cryo-EM micrographs of RBD-ACE2 complex on RGO-supported region (a) and RGO-broken region (b). These two micrographs were taken from different regions of the same EM grid.
